# Phasic signaling in the bed nucleus of the stria terminalis during fear learning predicts within- and across-session cued fear expression

**DOI:** 10.1101/768416

**Authors:** Max Bjorni, Natalie G. Rovero, Elissa R. Yang, Andrew Holmes, Lindsay R. Halladay

## Abstract

While results from many past studies have implicated the bed nucleus of the stria terminalis (BNST) in mediating the expression of sustained negative affect, recent studies have highlighted a more complex role for BNST that includes aspects of fear learning in addition to defensive responding. As BNST is thought to encode ambiguous or unpredictable threat, it seems plausible that it may be involved in encoding early cued fear learning, especially immediately following a first tone-shock pairing when the CS-US contingency is not fully apparent. To investigate this, we conducted in vivo electrophysiological recording studies to examine neural dynamics of BNST units during cued fear acquisition and recall. We identified two functionally distinct subpopulations of BNST neurons that encode the intertrial interval (ITI) and seem to contribute to within- and across-session fear learning. “Ramping” cell activity during cued fear acquisition parallels the increase in freezing expression as mice learn the CS-US contingency, while “Phasic” cells encode post-shock (US_post_) periods (30 s following encounter with footshock) only during early trials. Importantly, the magnitude of Phasic unit responsivity to the first US_post_ period predicted not only freezing expression in response to the subsequent CS during acquisition, but also CS freezing evoked 24 hr later during CS retrieval. These findings suggest for the first time that BNST activity may serve as an instructive signal during cued fear learning.

## Introduction

The ability to recognize threats and execute appropriate defensive responses is crucial for survival. Many decades of research have identified key players in the acquisition, expression, and extinction of fear, including the amygdala, hippocampus, and prefrontal cortex (Fendt and Fanselow 1999, LeDoux 2000, Maren 2001, Orsini and Maren 2012). Recent work has also focused on the bed nucleus of the stria terminalis (BNST) and its involvement in sustained negative affect. Currently, the role of the BNST in aversive learning has not been fully defined, but recent studies (see Goode and Maren 2017 for review) have highlighted a complex role for BNST in fear learning and defensive responding. Uncovering intricacies of its role in aversive learning will be crucial for development of improved treatments for neuropsychological disorders related to negative affect.

Early work laid out a selective role for BNST in processing anxiety rather than fear; whereas explicit fear cues activate the amygdala, evidence suggested the BNST was important for processing information of longer duration and less specificity (Davis 1998, LeDoux et al. 1988). Two subsequent decades of work have improved this perhaps oversimplified distinction between amygdala versus BNST mediating fear versus anxiety. The BNST has since been identified as an important mediator of the limbic forebrain processes that encode aversive motivational states, specifically ambiguous or temporally unpredictable aversive cues (Davis et al. 2010, Daldrup et al. 2016, Goode and Maren, 2017, Naaz et al. 2019, Goode et al. 2019).

Past studies have reported that BNST is necessary for fear learning involving long-duration, unpredictable, or contextual stimuli, but not for shorter-duration or explicit fear learning (LeDoux et al. 1988, Sullivan et al. 2004, Waddell et al. 2006). Notwithstanding this, more recent *in vivo* electrophysiological recording studies have revealed that some BNST neurons encode conditioned stimulus (CS) onset (Haufler et al. 2013), as well as fear expression during both contextual and cued fear recall (Marcinkiewcz et al. 2016). Importantly, our recent electrophysiological evidence showed that in the same animals, BNST neurons encoded fear expression as well as encounter with footshock (Marcinkiewcz et al. 2016), suggesting a dual role for BNST – not only predicting potential aversive encounters, but potentially also encoding sensory feedback following such an encounter. This raises the question of whether dynamic neural changes in BNST contribute to the fear learning process despite BNST not being *necessary* for cued fear learning.

Present understanding of the seeming unimportance for BNST in some types of aversive learning is in large part due to previous tests of necessity that employed lesions or reversible inactivation (e.g., Gewirtz et al. 1998, Hammack et al. 2004, Sullivan et al. 2004, Waddell et al. 2006, Duvarci et al. 2009, Goode et al. 2015). With regard to fear learning, the brain is equipped with compensatory circuits that serve as secondary mechanisms for fear acquisition after damage to the “primary” fear circuit (Poulos et al. 2010, Zelikowsky et al. 2013). For example, BNST mediates fear learning following amygdala lesions (Poulos et al. 2010, but see Zimmerman and Maren 2011). As such, it seems plausible that the BNST encodes aspects of aversive learning in parallel with the amygdala. That BNST neurons encode both behavioral expression and sensory input during fear acquisition (Marcinkiewcz et al. 2016) seems to support this hypothesis. But uncovering the extent to which BNST contributes to aversive learning alongside parallel circuits is not fully possible using lesions or inactivations which lack temporal precision and cannot easily parse primary versus compensatory circuitry. To this end, we used *in vivo* electrophysiology to observe the neural dynamics of the non-manipulated BNST during cued fear acquisition, as well as both cued and contextual fear retrieval. Recordings revealed dynamic changes in the activity of two functional populations of BNST neurons that coincided with (and in some cases, predicted) both within- and across-session fear learning. These findings suggest for the first time that BNST activity may serve as an instructive signal during early fear learning.

## Results

### Behavior during fear acquisition and retrieval

On day 1, mice (n=10) acquired a fear response to the tone following 5 trials of CS + US pairings in Context A. Fear learning resulted in significantly different levels of freezing expression across trials (one-way ANOVA; F(5,45)=23.7, p<.0001; Figure 1A-B). Relative to baseline, freezing during the CS was significantly elevated during trials 4 and 5 (Tukey’s HSD, p<.0001). Auditory fear memory was retrieved on day 2, when mice received 10 CS-only presentations in Context B. CS-evoked freezing significantly decreased across the session (F(10,90)=2.717, p=.006), where average freezing during late trials (i.e., CS 6-10) was significantly attenuated compared to freezing during early trials (i.e., CS 1-5) (t(9)=2.88, p=.018; Figure 1C-D). To assess contextual fear memory, on day 3 mice were returned to Context A and were allowed to freely behave for 5 min in the absence of any CS or US presentations. Freezing was slightly elevated during minute 3 relative to the other time blocks (F(4,36)=5.174, p=.002, Tukey’s HSD p<.01; Figure 1D-E). Average time spent freezing during the CX test was similar to the levels seen on day 2 during CS retrieval, indicating that mice had acquired a fear response to both the discrete and contextual cues associated with fear learning.

**Figure 1.**
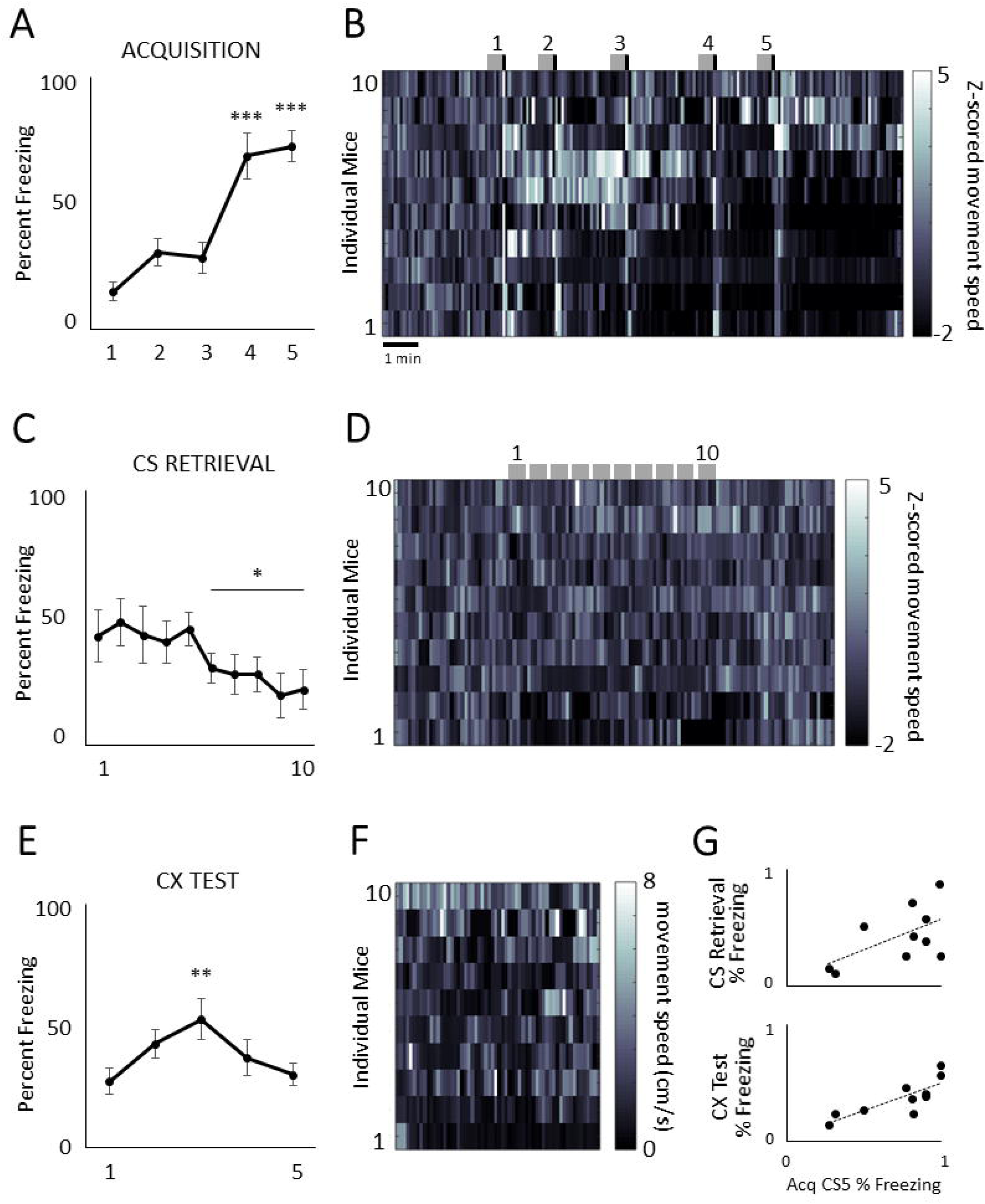
Freezing expression during cued fear acquisition, CS retrieval, and CX test. A) Mice (n=10) were trained to fear an auditory stimulus by pairing it with footshock in Context A. Freezing expression increased over the 5 acquisition trials, and was significantly different from baseline on trials 4 and 5. B) Individual animals’ movement speeds (normalized to baseline speeds) reflected US-induced movement and a reduction in movement during the CS and ITI over the course of the session. Gray boxes represent CS-on times, and black vertical bars indicate time of US delivery. C) On day 2, mice received 10 CS presentations in Context B. Relative to early trials (i.e., 1-5), CS-evoked freezing expression was reduced during late trials (i.e., 6 −10). D) Individual animals’ movement speed during the CS retrieval session, each normalized to its baseline movement speed. Gray boxes indicate CS-on times. E) On day 3, mice returned to Context A for 5 min to assess contextual fear expression. Freezing was significantly greater during block 3 versus other time blocks. F) Individual animals’ raw movement speed over the context exposure. G) Each animal’s freezing to the final CS on acquisition day positively correlated with freezing expression during CS retrieval (top) and CX test (bottom). Data in A, C, and E represent the mean ± S.E.M. * = p <.05, ** = p<.01, *** = p<.001.

Freezing expression during the final CS on acquisition day served as a reliable measure of fear learning for individual mice because it predicted freezing expression on subsequent test days (Figure 1G). There was a strong trend for a positive correlation between acquisition-day CS 5 freezing and freezing during the CS on CS retrieval (r=.593, p=.071). There was a significant positive correlation between acquisition-day CS5 freezing and freezing to the CX on day 3 (r=.81, p=.005). As such, the degree to which each animal froze to the final CS during acquisition was predictive of fear expression in subsequent tests of both the discrete and contextual fear memories.

### Classification of BNST cell types

Single units were classified based on their mean firing rate change across the acquisition session. For acquisition and CS retrieval sessions, the firing rate of each unit recorded was Z score normalized against its average firing rate during the 3 min baseline period of the session. For CX test, unit firing rates were normalized to a 3 min homecage period immediately prior to the start of the session. Spike data for each recording session were used to generate a perievent time histogram (PETH; bin size = 10 sec) aligned to session start for each unit. Bins with a Z score of >2.58 or <-2.58 were considered to significantly different than baseline. Inclusion of bins for each analysis are noted below.

Initial visual inspection of individual PETHs for acquisition (e.g., Figure 2A) indicated consistent firing rate changes during the ITI (i.e., the time bins between US offset and subsequent CS onset) for many units, as opposed to bins occurring during CS-on periods. In fact, only 3 units out of the 79 recorded (3.8%) significantly changed their firing rate during the CS and not the ITI (Figure 2D), indicating that very few BNST neurons solely encoded the presence of a discrete fear stimulus. As such, Z scores from the 6 time bins immediately following offset of the US (i.e., first min of each ITI) were used to classify cells into three types – Phasic, Ramping, and no response (Figure 2B-F).

**Figure 2.**
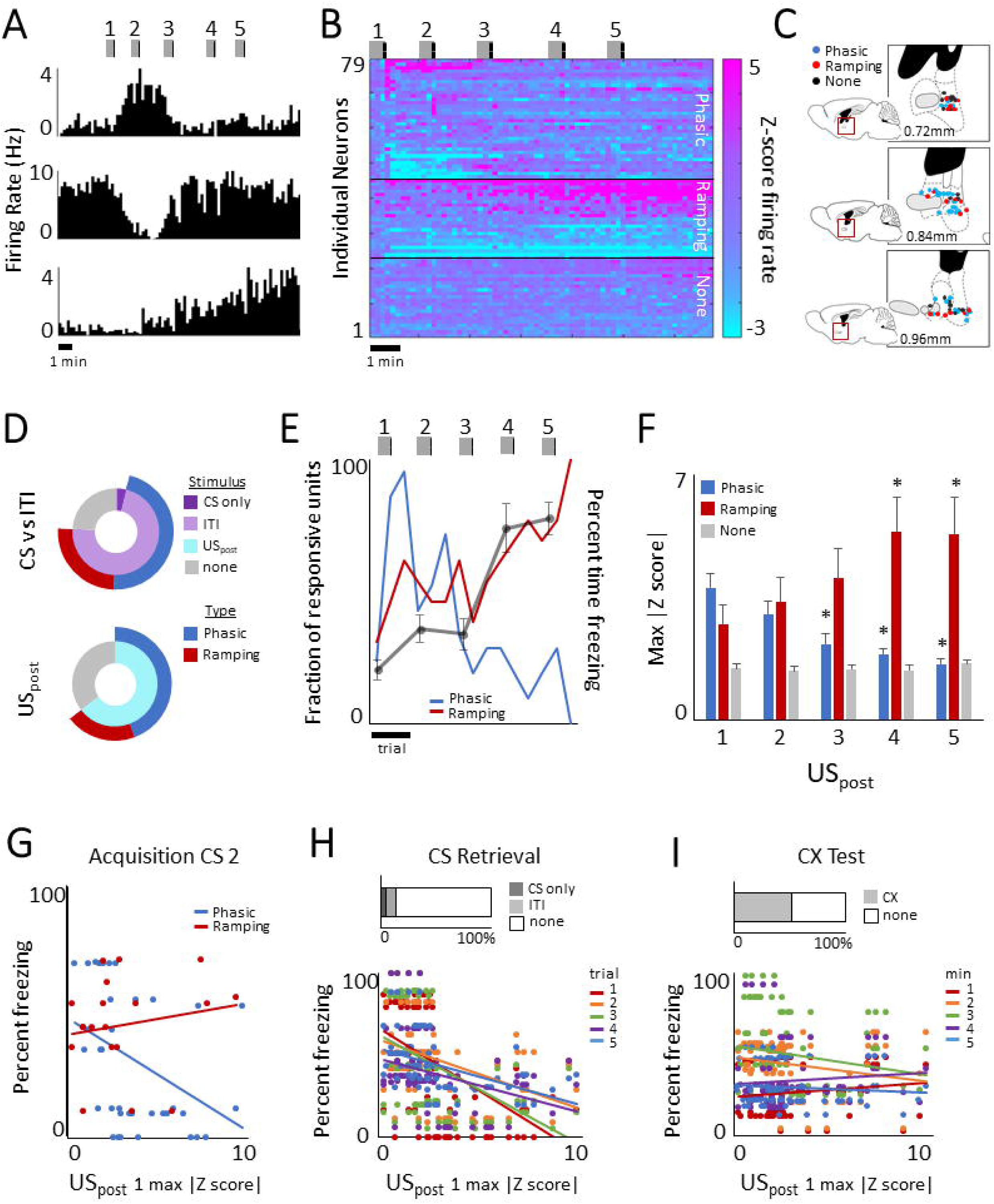
BNST phasic encoding predicts within- and across session fear expression. A) Perievent time histograms (PETHs; 10 sec bins) of representative BNST units recorded during the cued fear acquisition session. Phasic units (n=34 of 79) exhibited significant firing rate changes following early CS-US pairings that returned to baseline by later trials (top, middle), while Ramping units (n=19 of 79) exhibited gradual, sustained firing rate changes that emerged some time after the first shock presentation and persisted throughout the session (bottom). B) Population heat plot of Z scored activity (relative to 3 min baseline) of all units recorded across the acquisition session (10 sec bins, organized from greatest to least Z scored US_post_ response for each unit classification type). Phasic units exhibited significant changes in spike rate following US_post_ 1 or 2, while Ramping units exhibited significant rate changes lasting through the end of the session. C) Histological reconstruction of recorded units, color coded by unit type. Blue = Phasic, red = Ramping, and black = no response. Placements overlaid on sagittal atlas sections, 0.72, 0.84, and 0.96 mm lateral relative to midline. D) The majority of units recorded during acquisition were responsive to the ITI (top, lilac) as well as the US_post_ periods (bottom, aqua). Very few units were responsive only to the CS-on period (top, violet). ITI and US_post_ responsive units were comprised of both Phasic (blue) and Ramping (red) units. E) Average freezing behavior (gray) overlaid on line graphs depicting the fraction of responsive Phasic (blue) and Ramping (red) units for the CS-on, US_post_, and ITI periods across trials. As freezing expression increased across trials, fraction of responsive Phasic units decreased as the fraction of responsive Ramping units increased. F) For each unit type, average maximum absolute Z score during the US_post_ period for each trial. Average Phasic response to the US_post_ decreased across trials, while average Ramping response to the US_post_ increased across trials. G) Responsiveness to the first US_post_ period negatively predicted freezing behavior during the subsequent CS presentation for Phasic, but not Ramping, units. H) On day 2, very few of the 81 units recorded were responsive to either the CS-only or ITI periods (top). However, acquisition day responsiveness to the first US_post_ period negatively predicted CS-evoked freezing behavior during early trials in Context B 24 hrs later. I) About half of all units recorded on day 3 (n=74) when mice returned to Context A were responsive to the context. Unlike previous days, responsiveness to the first US_post_ period on acquisition day did not predict freezing to the context on day 3. Panels A, B, and E: Gray boxes represent CS-on times, and black vertical bars indicate time of US delivery. Data in E and F represent the mean ± S.E.M. * = p <.05.

A large proportion of units recorded (43%, 34 of 79) exhibited phasic significant firing rate changes during at least one of the first 3 ITIs, but specifically returned to baseline firing rate by ITI 4, and thus were classified as Phasic units (e.g., Figure 2A top, middle). About one fourth of recorded units (24%, 19 of 79) exhibited significant firing rate changes during or after the first ITI that persisted, and in many cases increased in magnitude or duration, throughout the rest of the session (e.g., Figure 2A bottom). As such, these units were classified as Ramping units. Units that did not exhibit significant firing rate changes during the ITIs were considered to have no response (33%). Figure 2E depicts the change in fraction of responsive units by type, with average percent time freezing during the CS overlaid; Phasic units exhibit significant firing rate changes prior to the emergence of CS-evoked freezing, indicating that the units may encode uncertainty or initial learning. On the other hand, Ramping units primarily exhibit significant rate changes following an increase in CS-evoked freezing, suggesting that they may encode certainty or late learning.

### Neural responses to the US_post_ and ITI periods

Although accurate recording during the footshock is not possible due to electrical interference from the grid floor occluding the signal during the 2 sec shock, we observed prolonged responses immediately following shock delivery, in some cases lasting upwards of 20 sec in both Phasic (94.6%) and Ramping (80%) cells (Figure 2D). Using each unit’s maximum absolute Z score of the three bins (i.e., 30 sec window) following each shock (US_post_), a 3×5 mixed-design ANOVA revealed an interaction between cell type and trial (F(8,300) = 7.793, p<.00001), whereby on average, Phasic cells showed a population-wide gradual decrease in their magnitude of response to the US_post_ across the session (Figure 2F); post hoc analysis revealed that response magnitude differed significantly on trials 3-5 versus trial 1 (p<.05). Alternatively, Ramping cells exhibited a population-wide increase in magnitude of response across the session, with peak magnitude following US 4; post hoc tests showed that trials 4 and 5 differed significantly compared to trial 1, (p<.05). No Response cells did not show any significant population encoding of the US across the session.

ITI and US_post_ responsiveness suggested encoding of learning by some BNST units. As such, to determine whether shock response magnitude was an indication of fear learning, for units that were responsive to the US (47 of 79) we used simple linear regression to compare the magnitude of each unit’s response to the US_post_ period (i.e., the maximum z score of the 3 bins following each shock) with freezing during the subsequent CS (Figure 2G). We found that Phasic cell encoding during the post-shock period following US 1 significantly predicted percent time spent freezing during CS 2 (F(1,35) = 6.081, p=.019, r = -.390) such that the greater the firing rate during the post shock period, the less the freezing during the subsequent CS. US_post_ responsiveness did not predict CS freezing in Ramping units (F(1,19) = .530, p=.476, r = .169) or in Phasic units on any other trial. Results suggest that Phasic units may primarily encode attention or uncertainty of aversive contingency, and thus are negatively correlated with early fear learning.

### Neural responses during cued fear retrieval

Neurons recorded during CS retrieval (n=81) were analyzed to determine whether firing rate changes over the session reflected similarities with the Phasic and/or Ramping cells recorded during acquisition. To determine stimuli responsiveness as learning took place across the session, firing rates were analyzed separately for early (i.e., 1-5) and late (i.e., 6-10) trials. Similar to acquisition, recording data for CS retrieval also took into consideration the ITI period in addition to CS presentations. A unit was considered to be CS-only responsive if both its Z-scored firing rate for any 10 sec bin during any CS window had a Z score of >2.58 or < −2.58, and its firing rate Z-score during bins of the ITI period (one 10 sec bin per ITI) did *not* exceed ±2.58. BNST was minimally responsive to CS-only presentations; only 3.7% (3 of 81 units) were responsive to early CS presentations (Figure 2H top), and 4.9% (4 of 81) during late CS presentations. No units were responsive to only the CS during both early and late trials. A slightly greater proportion of units was responsive during the ITI periods; 9.9% (8 of 81) of units in early trials and 16% (13 of 81) of units in late trials exhibited significantly altered firing rates during the ITI period. Similar to the CS-only responsive cells, units responsive during the ITI rarely responded during both early and late trials (3 of 81), suggesting that BNST neurons primarily encoded phasic changes during the CS retrieval session as opposed to sustained activity seen in Ramping units the day prior.

Next, we investigated whether performance on fear retrieval could be predicted based on population-wide BNST encoding during fear acquisition, since neural magnitude of response to the first shock on acquisition day predicted within session freezing. Indeed, the magnitude of individual units’ responsiveness to the initial shock during acquisition correlated significantly with freezing expression during each of the 5 early CS retrieval trials (r = -.514, -.386, -.451, -.323, -.462, p<.001), such that the greater the magnitude of response to the post shock period, the less CS-evoked freezing the animal displayed (Figure 2H bottom). Similar to acquisition, this suggests that greater BNST encoding following the first tone-shock pairing predicted less overall fear learning (as indexed by freezing expression) in addition to within session learning, which could reflect aversive outcome uncertainty.

### Neural responses during context fear retrieval

Mice were returned to the acquisition context on day 3 to assess contextual fear learning; as there was no baseline period to analyze, spikes recorded during exposure to the context were binned into 1-min blocks and normalized to each unit’s homecage firing rate (i.e., 3 min recording occurring just prior to the animal being placed into the acquisition context). Similar to above, units were considered to be responsive to the context if their Z score for that block exceeded ±2.58. We found 52% of units recorded (28 of 74) to significantly change their firing rate after returning to the acquisition context (Figure 2I top) during at least one of the 1-min blocks. Unlike acquisition and CS retrieval, however, unit responsiveness to the first shock pairing on acquisition day did not predict freezing to the context on day 3 during any time block (r=.13, -.221, -.179, .09, .08, p>.05; Figure 2I bottom).

## Discussion

Electrophysiological recordings of BNST neurons during fear acquisition revealed two subpopulations responsive to the task: Ramping cells gradually exhibited firing rate changes that persisted throughout the rest of the session; the fraction of Ramping units responsive to the US_post_ and ITI phases increased in parallel with magnitude of freezing expression as mice learned the CS-US contingency. Conversely, Phasic units exhibited significant firing rate changes following the initial tone-shock pairings that returned to baseline before the last trials. This Phasic cell encoding predicted both within- and across-session learning because Phasic units’ magnitude of responsiveness following the first shock presentation negatively correlated with freezing expression during the subsequent CS, as well as freezing during early CS presentations 24 hours later during cued fear retrieval. These findings suggest that Phasic BNST activity during non-CS periods of cued fear learning may contribute to the acquisition and retention of cued fear.

BNST Phasic neurons maximally encoded the US_post_ period following the unexpected delivery of shock in early acquisition trials. These data echo previous electrophysiological findings that components of neural fear circuitry (i.e., lateral amygdala (LA) and periaqueductal gray (PAG) neurons) preferentially encode unsignaled versus signaled shock, and consequently, neural encoding decreases across fear acquisition sessions (Johansen et al. 2010). Connections between PAG and amygdala modulate fear learning and expression based on instructive signaling and feedback (Johansen et al. 2010, Ozawa et al. 2017). It has been proposed that PAG may indirectly convey nociceptive information related to shock to the LA to enable plasticity and induce fear learning (Kim et al. 2013, Herry and Johansen 2014). But since PAG does not directly project to LA, regions such as thalamic nuclei or anterior cingulate cortex have been proposed as mediators (Shi and Davis 1999, Lanuza et al. 2004, Tang et al. 2005, McNally et al. 2011, Herry and Johansen 2014). However, based upon Phasic US_post_ signaling recorded here, we propose BNST as an additional potential instructor of fear learning. BNST has been shown to encode affective aspects of pain (Deyama et al. 2007, Morano et al. 2008, Minami and Ide 2015), and it is well-situated to relay information to the amygdala; PAG mediates nociception via projections to BNST (Li et al. 2016), and in turn, BNST projects to subregions of the amygdala including LA, basal (BA) and central (CeA) nuclei (Dong et al. 2000, Gungor et al. 2015, Krüger et al. 2015). As such, BNST signaling may relay information related to noxious events to inform future defensive response selection. This is evidenced not only by the ability of the magnitude of responsiveness following the first shock to predict CS-evoked freezing behavior on the subsequent trial, but also its ability to predict cued freezing expression 24 hours later during CS retrieval.

While our findings suggest a key mediating contribution of the BNST for fear learning, this role does not appear to be necessary for such learning under normal circumstances. Several past studies have shown that BNST is not necessary for some types of fear learning (Gewirtz et al. 1998, Hammack et al. 2004, Sullivan et al. 2004, Waddell et al. 2006, Duvarci et al. 2009). However, the importance of the BNST can be unmasked following damage to the basolateral (BLA) amygdala (Poulos et al. 2010). Poulos and colleagues (2010) showed that extended fear conditioning (i.e., >5 trials) can result in within- and across-session freezing following damage to the BLA, so long as the BNST was intact. Importantly though, lesions to either BLA *or* BNST attenuated within-session learning during the initial trials (i.e., 1-4), and BNST lesions *alone* also impaired fear recall 24 hours later. These results suggest that BNST does in fact play a role in fear learning, both during initial US occurrences as well as long-term retention. Phasic signaling in BNST reported here, which predicted both within- and across-session learning supports this idea. Furthermore, the parallel increase in freezing and Ramping cell responsiveness across acquisition also indicates potential involvement of Ramping signaling in within-session learning.

One caveat, however, is that compensatory BNST fear learning has been demonstrated in contextual (Poulos et al. 2010) but not cued fear conditioning (Zimmerman and Maren 2011). Although evidence here supports a role for BNST involvement in cued fear learning, we like others did not find sufficient evidence to conclude that BNST encodes fear cues specifically. Rather, responsive neurons exhibited firing rate changes during the US_post_ and ITI periods when stimuli present included only the context. Several lines of evidence have shown that BNST encodes contextual fear conditioning (Sullivan et al. 2004, Waddell et al. 2006, Zimmerman and Maren 2011, Davis and Walker 2014, Hammack et al. 2015). During early cued fear learning, some contextual conditioning takes place because the associative strength of the US has not been fully attributed to the CS (Rescorla et al. 1972). Thus, for Phasic BNST neurons specifically, it is possible that responsiveness during non-CS periods is indicative of some degree of contextual fear learning, which subsides after repeated CS-US pairings as the majority of associative strength shifts to the CS.

Past *in vivo* electrophysiology studies have shown that LA and PAG firing rate changes in response to fear cues often correlate with fear expression in real-time (Blair et al. 2003, Johansen et al. 2010, Halladay and Blair 2015). Here, subpopulations differ in this regard since Phasic signaling inversely predicted subsequent cue-evoked fear expression, but Ramping cell responsiveness and freezing increased in parallel across the session. Some recent recording studies have identified opposing BNST populations that distinctly encode either anxiogenic or anxiolytic behaviors (Jennings et al. 2013, Kim et al. 2013). While our data do not suggest Phasic and Ramping cells oppose each other directly, the differences in neural response patterns highlight the heterogeneity of neural activity in BNST previously reported (Gungor and Paré 2016). One important distinction here versus past recording studies is that Phasic ITI signaling predicted behavior as far as 30-90 sec in the future (i.e., during CS presentation), rather than coinciding with concurrent behavioral expression. Similarly, while Ramping cell responsivity mirrored increase in freezing behavior over time, we did not find evidence that Ramping cells directly modulated behavior in real time. This seems to reflect sustained encoding of aversive information, in support of previous studies implicating BNST in modulating responses during sustained or long-duration aversive states (Waddell et al. 2006, Walker and Davis 2008, Davis et al. 2010).

Finally, it is worth noting that the increase in Ramping cell responsiveness observed across the acquisition session is reminiscent of past studies showing an increase in CeA neural responsiveness to the CS across fear learning as freezing increases (Duvarci et al. 2011, Ciocchi et al. 2010). As BNST-projecting CeA corticotropin-releasing factor (CRF) neurons are necessary for sustained fear expression (Asok et al. 2018), it seems plausible that BNST Ramping cells may receive input from CeA that drives the long-duration ITI responsiveness reported here. This would be an interesting direction for future circuit-specific studies.

In summary, we identified two functionally discrete subpopulations of BNST neurons that encode the US_post_ and ITI periods of cued fear learning. Phasic signaling in BNST may instruct both within- and across-session fear expression by relaying information specific not to the CS, but rather, to the longer duration periods between acquisition trials. Future studies such as those including multi-site *in vivo* recordings will be necessary to more fully understand the implications of phasic neural signaling in BNST during cued fear learning.

## Materials and Methods

### Subjects

Male C57BL/6J mice were obtained from The Jackson Laboratory (Bar Harbor, ME; https://www.jax.org/strain/000664) at 7-8 weeks of age and housed in pairs in a temperature (72 ± 5 °F) and humidity (45 ± 15%) controlled vivarium, under a 12 hr light/dark cycle (lights on at 0630 hr). All procedures were approved by the NIAAA Animal Care and Use Committee and followed the NIH guidelines outlined in *Using Animals in Intramural Research*, as well as the local Animal Care and Use Committees.

### Behavioral procedures

Behavioral sessions were conducted using MedPC (Med Associates, Fairfax, VT) in conjunction with Plexon recording equipment described below. Acquisition took place on day 1 in Context A, 27 × 27 × 11 cm chamber with a metal rod floor for shock delivery, cleaned with a 69% water/30% ethanol/1% vanilla-extract solution. Acquisition consisted of a 180-sec baseline period followed by 5 pairings of a pure tone CS (30 sec, 3 kHz, 75 dB) co-terminating with a footshock US (2 sec, 0.6 mA), with variable ITIs between 60-120 sec. Mice were tested for cue-evoked freezing during CS retrieval 24 hr later, which took place in Context B, 20 cm diameter Plexiglas cylinder with a solid, opaque floor and cleaned with a 99% water/1% acetic acid solution. The session consisted of a 180-sec baseline period followed by 10 CS presentations (10 sec ITI). 24 hr following retrieval, mice were returned to Context A for a 5-min context (CX) test. Freezing behavior was scored using the automated freezing function in CinePlex Editor (Plexon Inc., Dallas, TX) and verified by hand scoring by blind experimenters. We used the automatically scored freezing intervals for analyses due to the high temporal resolution required for spike timing data.

### *In vivo* neuronal recordings

At least 1 week prior to experimentation, mice (n=10) were surgically implanted with fixed microelectrode arrays (Innovative Neurophysiology, Durham, NC) containing 2 rows of 8×35 µm tungsten electrodes, with 150 µm electrode-spacing and 200 µm row spacing, unilaterally implanted in BNST (array center at 0.3 mm anterior, 0.8 mm lateral, and 4.15 mm ventral relative to Bregma). Electrophysiological recordings were conducted during acquisition, CS retrieval, and CX test sessions, using the OmniPlex D Neural Data Acquisition System, with simultaneous behavioral analysis via CinePlex Behavioral Research Systems (Plexon Inc., Dallas, TX). Waveforms exceeding a set voltage threshold were digitized at 40 kHz. Manual cluster analysis and inspection of waveforms and inter-spike intervals were performed offline using Plexon Offline Sorter. Spike and timestamp information were integrated and analyzed using NeuroExplorer (NexTechnologies, Littleton, MA).

Following completion of experiments, mice were anesthetized with 2% Isoflurane (in oxygen, 2L/min), and a current stimulator (S48 Square Pulse Stimulator, Grass Technologies, West Warwick, RI) was used to deliver 2 sec of 40 µA current through each electrode to make a small marking lesion. 24 hr later, mice were overdosed with ketamine/xylazine and perfused transcardially with 4% paraformaldehyde in phosphate buffered saline. Brains were sectioned on a vibratome (Leica VT100S) and 50 µm sections slide-mounted and stained for histological verification of electrode placements. Only units from electrodes confirmed to be located within BNST were included in the analyses.

### Experimental Design and Statistical Analyses

All animal and unit n values are described for each of the three behavioral sessions in Results below. Behavior data were analyzed using One- or Two-Way Analysis of Variance (ANOVA) and Tukey’s Honest Significant Difference (HSD) test for planned post hoc comparisons. For comparison of behavior during early versus late trials on day 2, data were analyzed using paired sample Student’s *t*-test. Results are presented as the mean ± S.E.M.

Spike data were normalized and analyzed as previously described (Halladay and Blair 2015, Gamble-George et al. 2016, Marcinkiewcz et al. 2016, Gunduz-Cinar et al. 2019, Hardaway et al. 2019). Classification of cell types based on responsiveness to events described below in Results. Comparisons between unit firing rates and behavior were made using simple linear regression or bivariate correlation.

